# O-GlcNAc transferase modulates formation of clathrin-coated pits

**DOI:** 10.1101/2022.06.17.496621

**Authors:** Sadia Rahmani, Hafsa Ahmed, Osemudiamen Ibazebo, Eden Fussner-Dupas, Warren W. Wakarchuk, Costin N. Antonescu

## Abstract

Clathrin-mediated endocytosis (CME) controls the internalization and function of a wide range of cell surface proteins. CME occurs by the assembly of clathrin and many other proteins on the inner leaflet of the plasma membrane into clathrin-coated pits (CCPs). These structures recruit specific membrane protein cargo destined for internalization and trigger the generation of membrane curvature that precedes eventual scission of CCPs from the plasma membrane to yield intracellular vesicles. The diversity of cell surface protein cargo thus controlled by CME indicates that CCP formation is regulated to allow cellular adaptation under different contexts. Of interest is how cues derived from cellular metabolism may regulate CME, given the reciprocal role of CME in controlling cellular metabolism. The modification of proteins with O-linked β-N-acetylglucosamine (O-GlcNAc) is sensitive to nutrient availability and may allow cellular adaptation to different metabolic conditions. We examined how the modification of proteins with O-GlcNAc may control CCP formation and thus CME. We used perturbation of key enzymes responsible for protein O-GlcNAc modification, as well as specific mutants of the endocytic regulator AAK1 predicted to be impaired for O-GlcNAc modification. We identify that CCP initiation and the assembly of clathrin and other proteins within CCPs is controlled by O-GlcNAc protein modification. This reveals a new dimension of regulation of CME and highlights the important reciprocal regulation of cellular metabolism and endocytosis.

## Introduction

The function of the many proteins at the cell surface is regulated by clathrin-mediated endocytosis (CME), the principal mechanism of internalization of integral membrane proteins at the plasma membrane (1–6). CME occurs as a result of the formation of clathrin-coated pits (CCPs) by the assembly of clathrin, the adaptor protein complex AP2, and many other proteins on the inner leaflet of the plasma membrane (3, 7–11). CCP formation is coupled to recruitment of specific integral membrane proteins (cargo) and generation of membrane curvature, leading to eventual scission from the plasma membrane to yield intracellular vesicles harboring protein cargo (1, 2).

The formation of CCPs and CME of specific proteins regulates the abundance and function of a wide range of cell surface membrane proteins. As such, this process is highly regulated by cues such as those derived from substrate stiffness (12), metabolism (13, 14), and growth factor stimulation (15). Thus, understanding the cues that regulate CCP formation and thus allow CME to gate the function of cell surface proteins is essential to understand cellular adaptation to changing conditions.

The initiation of CCPs is dependent on the recruitment of the central adaptor protein AP2 to the plasma membrane, which then supports clathrin recruitment. AP2 membrane recruitment is enhanced by binding to the FIE complex comprised of FCHo, Intersectin and Eps15 (16, 17) and by a conformational change in AP2 following binding to phosphatidylinositol-(4,5)-bisphosphate (PtdIns(4,5)P2) and specific sequence motifs on cargo proteins (5). The conformational change in AP2 is further stabilized by phosphorylation by AP2-associated kinase-1 (AAK1) or BMP2K on Thr156 of the µ2-subunit of AP2 (5, 18–22).

Following CCP initiation, clathrin, AP2 and many other proteins assemble into CCPs. While this assembly of proteins can lead to formation of clathrin-coated vesicles, a large proportion of nascent CCPs undergo abortive turnover at the plasma membrane without leading to formation of vesicles (5, 6, 21, 23). This indicates the existence of an endocytic checkpoint, in which the early stages of CCP formation undergo surveillance to support the further progression of suitable CCPs for vesicle formation, and the disassembly of other structures. The GTPase dynamin2 (6, 24, 25) and AAK1-mediated phosphorylation of AP2 µ2 (21) may regulate abortive turnover of some CCPs. However, how AAK1 is regulated to thus control CCP assembly remain poorly understood.

The generation of membrane curvature within CCPs is required for the eventual formation of internalized coated vesicles. Membrane curvature within CCPs can be driven by several proteins, including Epsin and CALM, or BAR-domain containing proteins such as endophilin, amphiphysin and SNX9 (1–3). CCPs may acquire curvature subsequent to assembly of clathrin and other proteins at the plasma membrane (26, 27), or throughout the assembly of CCPs (28). Both mechanisms can be resolved in the same cell (29), suggesting that CCP assembly and curvature generation can be flexible (30). However, not all clathrin assemblies acquire curvature, and the assembly of so-called flat clathrin plaques may be triggered by growth factor receptor activation and serve to control signaling outcome rather than mediate internalization (31).

The size of CCPs is also subject to regulation, which can occur for example as a result of recruitment of certain cargo proteins such low-density lipoprotein receptor as well as by the specific adaptors dab2 and ARH to CCPs (32). CCP size is also regulated by PtdIns(4,5)P2 levels (33) and specific CCP accessory proteins such as CALM and NECAP (34, 35). Notably, CALM regulates CCP size and curvature, suggesting that generation of membrane curvature and CCP are linked (35). Hence, both CCP size and membrane curvature can be modulated to control CCP function, but which specific cellular signals control CCP size and/or curvature remains incompletely understood.

Signals derived from cell metabolism are emerging regulators of endocytic membrane traffic (13, 14, 36). For example, the metabolic sensor AMP-activated protein kinase (AMPK) is activated under conditions of nutrient limitation. AMPK activation controls the endocytosis of specific proteins such as facilitative glucose transporters GLUT1 (37) and GLUT4 (38) and many others (13, 14). We previously found that many proteins including β1-integrin undergo changes in cell surface abundance upon AMPK activation (39). Collectively, these works suggest that metabolic signals may exert additional, still poorly appreciated effects on regulation of endocytic membrane traffic.

A regulatory mechanism that has emerging roles in linking cell metabolism with regulation of various cellular process is the dynamic modification of nucleocytoplasmic proteins with 2-acetamido-2-deoxy-D-glucopyranose (GlcNAc) linked by a β-glycosidic bond to serine or threonine residues of proteins (O-GlcNAc) (40–42). The dynamic addition and removal of O-GlcNAc from proteins is regulated by two enzymes: a glycosyltransferase (uridine diphosphate-*N*-acetylglucosamine:polypeptide β-acetylglucosaminyltransferase; OGT) (43) and a glycoside hydrolase (O-GlcNAcase; OGA (44). Deletion of OGT results in embryonic lethality, demonstrating the importance of this post-translational modification (45).

Unlike the various phosphatases and kinases that each control phosphorylation and dephosphorylation of a small number of specific proteins, O-GlcNAc cycling is controlled by a single glycosyltransferase, OGT, and has a single glycosidase, OGA. The single OGT and the single OGA can target as many as 4,000 proteins for O-GlcNAc modification (45, 46). Importantly, O-GlcNAc protein modification uses uridine diphosphate-*N*-acetylglucosamine (UDP-GlcNAc) as a substrate, a metabolite formed through hexosamine biosynthetic pathway (47). As such, the availability of specific nutrients such as glucose and glutamine modulate the extent of O-GlcNAc posttranslational modifications catalyzed by OGT and OGA, linking cell metabolism to control of a wide range of cellular processes (40–42).

O-GlcNAcylation impacts the function of target protein in many ways specific to the protein being modified, which includes changes in protein half-life, enzyme activity, subcellular localization and protein-protein interactions. Interestingly, O-GlcNAc modification of COPI and COPII components suggest regulation of biosynthetic membrane traffic (48–50). In addition, the key regulator of CCP assembly and function, AAK1, is subject to O-GlcNAc modification (51). However, how OGT and OGA, and in turn protein O-GlcNAcylation may regulate CCP formation and CME is not known. Here, we examine how perturbations of OGT and OGA modulate the initiation, assembly, maturation and scission of CCPs, and how dynamic protein O-GlcNAcylation of key CCP regulatory proteins such as AAK1 may impact this phenomenon.

## Results

### Protein O-GlcNAcylation regulates CCP dynamics

To examine how OGT and protein O-GlcNAcylation may regulate CCP formation and dynamics and thus CME, we used siRNA gene silencing. As expected, OGT silencing results in a robust reduction in both OGT expression as well as the levels of protein O-GlcNAc modification (**Figure 1A**). We then used this silencing of OGT in combination with total internal reflection fluorescence (TIRF) microscopy to obtain time-lapse image series of cells expressing eGFP-tagged clathrin light chain (4–6). In TIRF microscopy, the intensity of fluorophores and thus eGFP-clathrin objects declines with distance from the coverslip interface and therefore the acquisition of membrane curvature by CCPs. Hence, we performed imaging using near-simultaneous TIRF and widefield epifluorescence imaging modes (**Figure 1B**), as the latter does not exhibit intensity changes based on nanoscale changes in CCP curvature (6). To detect and analyze CCP dynamics, these image series are subject to automated image analysis of involving detection, tracking and analysis of diffraction-limited eGFP-clathrin objects using the TIRF channel for CCP detection. This approach was previously used to reveal how various perturbations or other experimental conditions affect specific stages of CCP initiation, clathrin assembly, curvature generation and CCP maturation (6).

**Figure 1.**
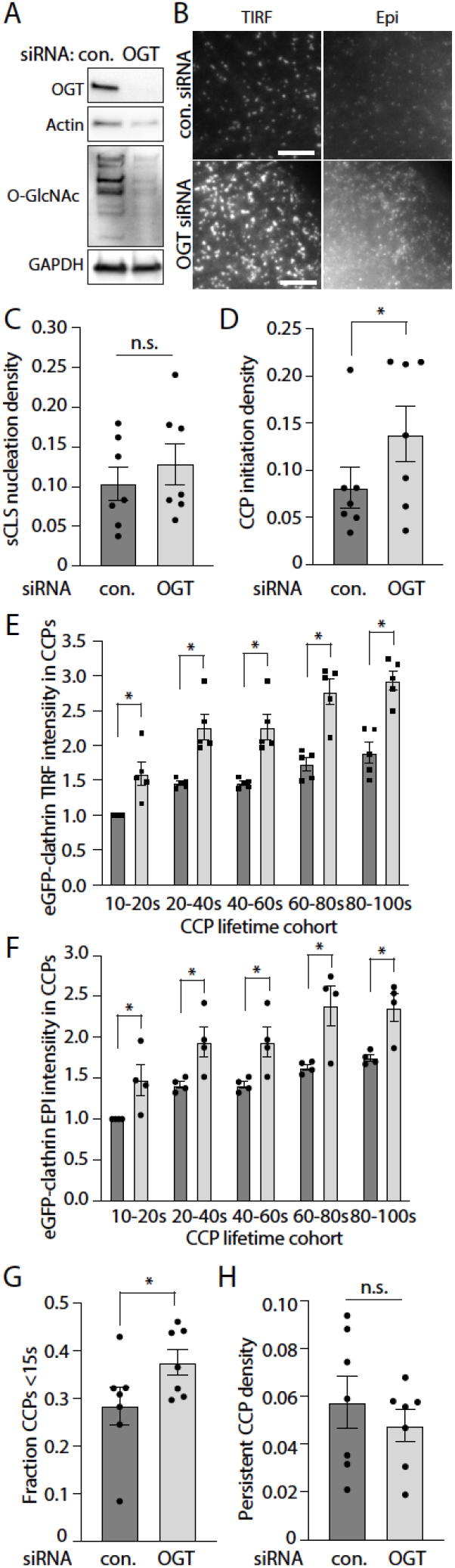
OGT silencing impacts CCP initiation, size, and lifetimes. ARPE-19 cells stably expressing eGFP-clathrin light chain were treated with siRNA to silence OGT or with non-targeting (control) siRNA. (A) Shown are representative immunoblots of whole cell lysates probed with antibodies as indicated. (B-H) Following transfection, cells were subjected to imaging with near-simultaneous time-lapse TIRF and epifluorescence microscopy. Shown in (B) are representative images, scale 5 µm. These time-lapse image series were subjected to automated detection, tracking, and analysis of CLSs as described in Materials and Methods. Shown are the initiation rates of sCLSs (C) and bona fide CCPs (D). Also shown are the intensities of eGFP-clathrin in CCPs based on lifetime cohorts detected in TIRF (E) or epifluorescence (F) time lapse image series, as well as the fraction of CCP lifetimes <15s (G) and the fraction of persistent CCPs (H). In each case (C-H), the data is presented as the mean (bar) ± SE from 7 independent experiments, as well as individual values from each experiment (dots). The number of total sCLS trajectories, CCP trajectories and cells (respectively) for each condition are: control siRNA 9987, 8733, 45 and OGT siRNA 13015, 15220, 42. *, p < 0.05

This method can resolve each clathrin object as either a *bona fide* CCP or a subthreshold clathrin labeled structure CLS (sCLS), as previously described (4–6). sCLSs are quantitatively and systematically distinguished from CCPs as they do not meet a minimum clathrin intensity threshold within the early stages of formation. As such, sCLSs likely represent stochastic assembly of clathrin at the plasma membrane that fail to proceed to formation of *bona fide* CCPs (4–6). Silencing OGT did not impact the rate of sCLS nucleation (**Figure 1C**), but in contrast resulted in a significant increase in the rate of initiation of *bona fide* CCPs (**Figure 1D**). This suggest that that loss of protein O-GlcNAcylation permits a more efficient assembly of proteins such as clathrin and AP2 at the earliest stages of CCP formation.

An increased efficiency of recruitment of clathrin and other proteins into CCPs at the earliest stages of CCP formation would be expected to lead to a higher rate or amount of eGFP-clathrin recruited per CCP. To test this, we examined the levels of eGFP-clathrin recruited to CCPs. Since the size of CCPs scales with CCP lifetime, we examine eGFP-clathrin intensity within CCP lifetime cohorts in each condition. Silencing OGT resulted in a robust and significant increase in the levels of eGFP-clathrin in clathrin structures in all lifetime cohorts in TIRF image series (**Figure 1E**). Since illumination of fluorophores in TIRF decays exponentially with distance from the coverslip and thus the cell surface, an increase in eGFP-clathrin intensity in TIRF images could reflect a reduction of membrane curvature generation and/or an increased recruitment of eGFP-clathrin to each object. This would in turn result in no change or increased eGFP-clathrin intensity in corresponding structures in widefield epifluorescence image series, respectively. OGT silencing robustly and significantly increased eGFP-clathrin intensity in clathrin objects in widefield epifluorescence image series (**Figure 1F**), suggesting that loss of OGT leads to enhanced recruitment of clathrin to CCPs and thus larger CCPs.

Following initiation, a subset of CCPs undergo abortive turnover, resulting in CCPs with shorter lifetimes (5, 6, 21, 23). To determine if the increased rate of eGFP-clathrin recruitment into CCPs upon OGT silencing was associated with changes in the rate of abortive CCP turnover, we examined CCP lifetimes. OGT silencing significantly increased in the fraction of CCPs with lifetimes <15s (**Figure 1G**) but did not change the proportion of CCPs that are persistent (present at the start and end of the time-lapse) (**Figure 1H**). These results suggest that while OGT silencing leads to an increased rate of CCP initiation and of eGFP-clathrin recruitment into nascent CCPs resulting in larger CCPs, these CCPs may have a higher propensity for abortive turnover.

Silencing OGT is an effective strategy to alter the levels of cellular protein O-GlcNAc modification, resulting in robust loss of O-GlcNAcylation (**Figure 1A**). We next sought to determine the impact of increasing cellular protein O-GlcNAcylation on CCP behaviour. However, silencing OGA also causes loss of OGT, which results in only a modest change in cellular protein O-GlcNAc modification upon OGA silencing (52). In contrast, treatment of cells with Thiamet G (TMG), a small molecule inhibitor of OGA, leads to a robust increase in cellular protein O-GlcNAc modification (**Figure 2A**). We thus examined how treatment with TMG impacts CCP dynamics as measured by near-simultaneous time-lapse imaging in TIRF and widefield epifluorescence microscopy (**Figure 2B**) and image analysis. TMG treatment did not alter the rate of sCLS nucleation (**Figure 2C**), or the rate of CCP initiation (**Figure 2D**).

**Figure 2.**
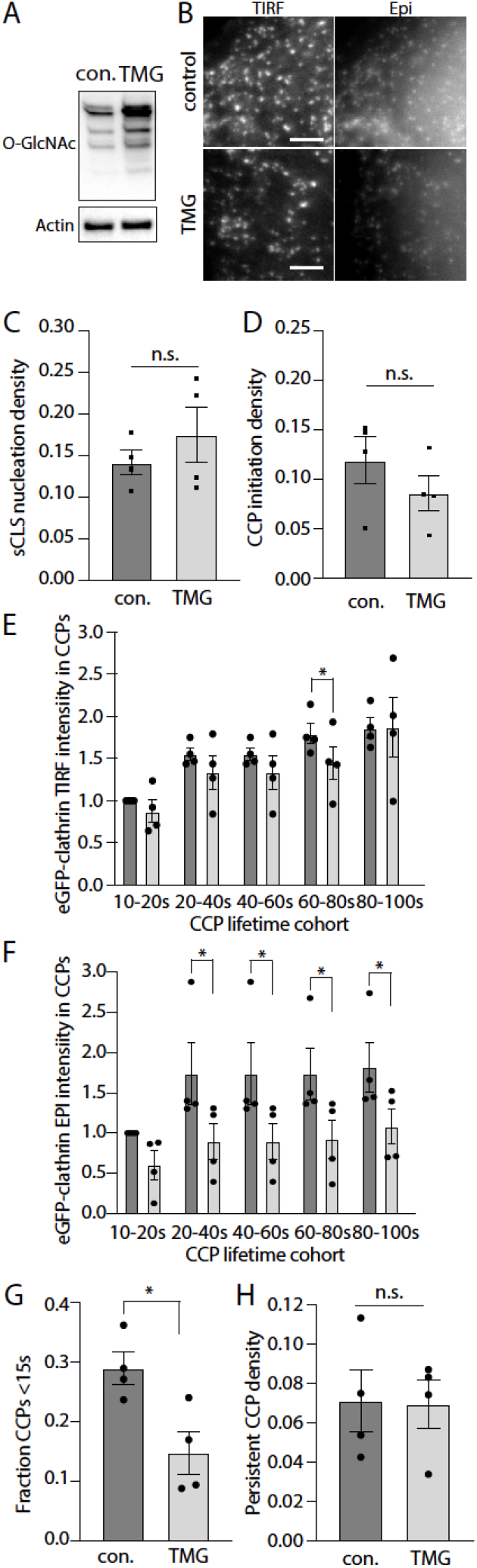
Inhibition of OGA with Thiamet G treatment impacts CCP size and lifetimes. ARPE-19 cells stably expressing eGFP-clathrin light chain were treated with 20 µM Thiamet G (TMG) or treated with vehicle control (con) for 4 hours. (A) Shown are representative immunoblots of whole cell lysates probed with antibodies as indicated. (B-H) Following transfection, cells were subjected to imaging with near-simultaneous time-lapse TIRF and epifluorescence microscopy. Shown in (B) are representative images, scale 3 µm. These time-lapse image series were subjected to automated detection, tracking, and analysis of CLSs as described in *Materials and Methods*. Shown are the initiation rates of sCLSs (C) and bona fide CCPs (D). Also shown are the intensities of eGFP-clathrin in CCPs based on lifetime cohorts detected in TIRF (E) or epifluorescence (F) time lapse image series, as well as the fraction of CCP lifetimes <15s (G) and the fraction of persistent CCPs (H). In each case (C-H), the data is presented as the mean (bar) ± SE from 4 independent experiments, as well as individual values from each experiment (dots). The number of total sCLS trajectories, CCP trajectories and cells (respectively) for each condition are: control 13063, 6479, 39 and TMG 10129, 3132, 21. *, p < 0.05

Our experiments with OGT silencing revealed that O-GlcNAc modification may suppress the rate of recruitment of eGFP-clathrin to CCPs and thus modulate CCP size. Consistent with this, TMG treatment resulted in a modest decrease in eGFP-clathrin in TIRF image series for some CCPs lifetime cohorts (**Figure 2E**), and a robust reduction in eGFP-clathrin per CCP in corresponding widefield epifluorescence image series (**Figure 2F**). The decrease in eGFP-clathrin per CCP in epifluorescence images indicates that TMG treatment reduces the levels of eGFP-clathrin recruited per CCP, while the modest effect on eGFP-clathrin levels per CCP in TIRF images upon TMG treatment may also indicate that CCPs formed under these conditions do not exhibit full curvature generation.

Interestingly, TMG treatment also robustly reduced the fraction of CCPs with lifetimes <15s (**Figure 2G**) and did not impact the fraction of persistent CCPs (**Figure 2H**), an observation complementary to the effect of OGT silencing. This suggests that TMG treatment reduces the abortive turnover of CCPs, favouring instead longer-lived productive CCPs. Collectively, thus in turn suggests that cellular protein O-GlcNAc modification may modulate the rate of recruitment of proteins into nascent CCPs, such that protein O-GlcNAcylation suppresses the rate of assembly of proteins such as eGFP-clathrin into CCPs. This in turn suggests that loss of O-GlcNAcylation may lead to aberrant formation or growth of CCPs, which elicits an increase in abortive CCPs.

### OGT controls cargo recruitment to CCPs

The increase in CCP size upon silencing OGT may reflect changes in CCP assembly and composition that also impact the recruitment of specific cargo proteins to CCPs. Given that some cargo such as Transferrin Receptor (TfR) and EGF Receptor (EGFR) are recruited to largely distinct CCPs and that CCPs harboring EGFR are uniquely regulated such as by Ca^2+^ signaling (15), we next examined how OGT and cellular protein O-GlcNAcylation may uniquely impact recruitment of EGFR and TfR to CCPs. To examine this, we labeled ARPE-19 cells that stably express eGFP-clathrin with A647-transferrin and rhodamine-EGF for 5 min, and subjected these to imaging by TIRF microscopy (**Figure 3A**), which we previously demonstrated to be a robust method for detecting changes in CCP cargo recruitment (15).

**Figure 3.**
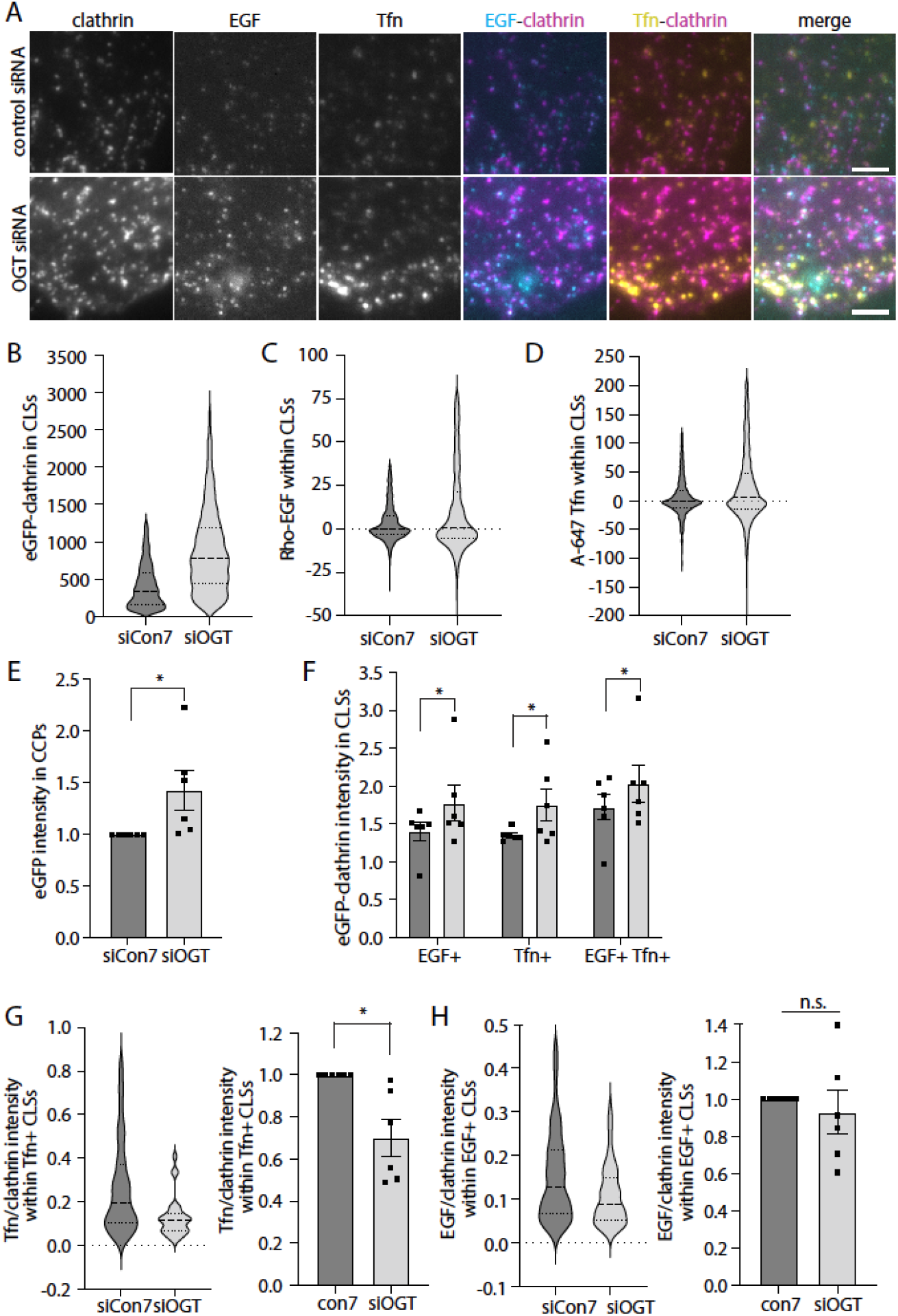
OGT silencing impacts cargo recruitment to CCPs. ARPE-19 cells stably expressing eGFP-clathrin light chain were treated with siRNA to silence OGT or with non-targeting (control) siRNA. Then, cells were treated with 20 ng/mL rhodamine-EGF (rho-EGF) and 10 μg/mL A647-Tfn for 5 min, followed by fixation and imaging by TIRF and epifluorescence microscopy, with representative images shown in (A), scale 3 µm. CLSs were subjected to automated detection and analysis as described in *Materials and Methods*. Shown are the distributions as a violin plot of measurements of eGFP-clathrin (B), rhodamine-EGF (C) and A647-Tfn (D) in individual CLSs within a representative experiment, as well as the mean (bar) ± SE of eGFP-clathrin in CLSs from 6 independent experiments, as well as individual values from each experiment (dots). (F-H) CLSs were sorted based on presence of EGF or Tfn as described in *Materials and Methods*. (F) Shown is the mean (bar) ± SE of eGFP-clathrin in CLSs sorted by cargo type from 6 independent experiments, as well as individual values from each experiment (dots). Also shown is the ratio of A647-Tfn/eGFP-clathrin (G) or rhodamine-EGF/eGFP-clathrin (H) intensities in CLSs, depicting the distribution of this ratio in a representative experiment as a violin plot (left panels), or the mean (bar) ± SE from 6 independent experiments, as well as individual values from each experiment (dots). The total number of CLSs quantified are as follows: control siRNA 18823 (total), 2826 (EGF+), 2828 (Tfn+); OGT siRNA 16227 (total), 2609 (EGF+), 2735 (Tfn+). *, p < 0.05

We examined these images through automated detection and analysis of clathrin-labelled structures (CLSs) using a Gaussian-based modeling approach (6), as was described in (15, 53, 54). We use the term CLS to denote clathrin structures detected in fixed samples, since identification of *bona fide* CCPs from short-lived subthreshold clathrin structures (sCLSs) requires live-cell analysis (5, 6). This approach allows study of the effect of OGT silencing on the recruitment of specific cargo to CLSs, as well as sorting of CLSs by cargo type to determine if there may be cargo-specific effects of protein O-GlcNAc modification on CCP formation and cargo recruitment.

As we observed in time-lapse image series, when examining individual CLSs, OGT silencing resulted in a robust increase in eGFP-clathrin in CLSs (**Figure 3B**), as well as an increase in A647-Tfn (**Figure 3C**), and rhodamine-EGF (**Figure 3D**) in these structures. As expected, eGFP-clathrin intensity levels within CLSs were significantly increased upon OGT silencing when considering multiple independent experiments (**Figure 3E**), and this effect was observed in all cohorts of CLS sorted by Tfn or EGF content (**Figure 3F**). This suggests that the increase in CCP size observed upon OGT silencing may not be specific to CCPs harboring EGFR or TfR.

The recruitment of specific receptors to CCPs upon OGT silencing may either scale with the size of CCPs, suggesting that OGT does not impact cargo recruitment separately from the control of CCP assembly, or may alter the proportion of receptor recruitment to clathrin structures. To distinguish between these possibilities, we examined the levels of A647-Tfn or rhodamine-EGF to individual CLSs. Interestingly, the ratio of A647-Tfn to eGFP-clathrin was reduced in OGT silenced cells, as seen in individual CLSs in a representative experiment (**Figure 3G,** *left panel*), as well in multiple independent experiments (**Figure 3G**, right panel). In contrast, the ratio of rhodamine-EGF to eGFP-clathrin was not impacted by OGT silencing (**Figure 3H**). This suggests that OGT silencing results in larger CCPs that proportionally recruit more EGFR yet are less effective at proportional recruitment of TfR.

### O-GlcNAc regulation of specific endocytic accessory proteins

Direct O-GlcNAc modification of specific proteins or specific residues has been difficult to resolve (55, 56). Methods to predict the O-GlcNAc modification of specific proteins recruited to CCPs are particularly useful. A recent comprehensive study that predicted O-GlcNAc modification of specific proteins revealed that many endocytic accessory proteins show likelihood of O-GlcNAc modification (57) (**Figure 4A**). Interestingly, proteins with significant intrinsically disordered regions such as AAK1, CALM/AP180 and Epsin are within this list, consistent with proteins with intrinsically disordered regions being preferred targets for O-GlcNAc modification (58). The possibility that the O-GlcNAc modification regulates AAK1, CALM and/or Epsin is consistent with central roles that these proteins play in formation of CCPs and agrees with the effects we observed upon disruption of cellular O-GlcNAcylation on CCP formation (**Figures 1-2**). Furthermore, AAK1 was previously shown to be modified by OGT in an OGT array, with 4 serine/threonine sites O-GlcNAc modified per AAK1, although the specific residues were not identified (51). Hence, we focused our efforts on elucidating the possible regulatory capacity of O-GlcNAc modification of AAK1.

**Figure 4.**
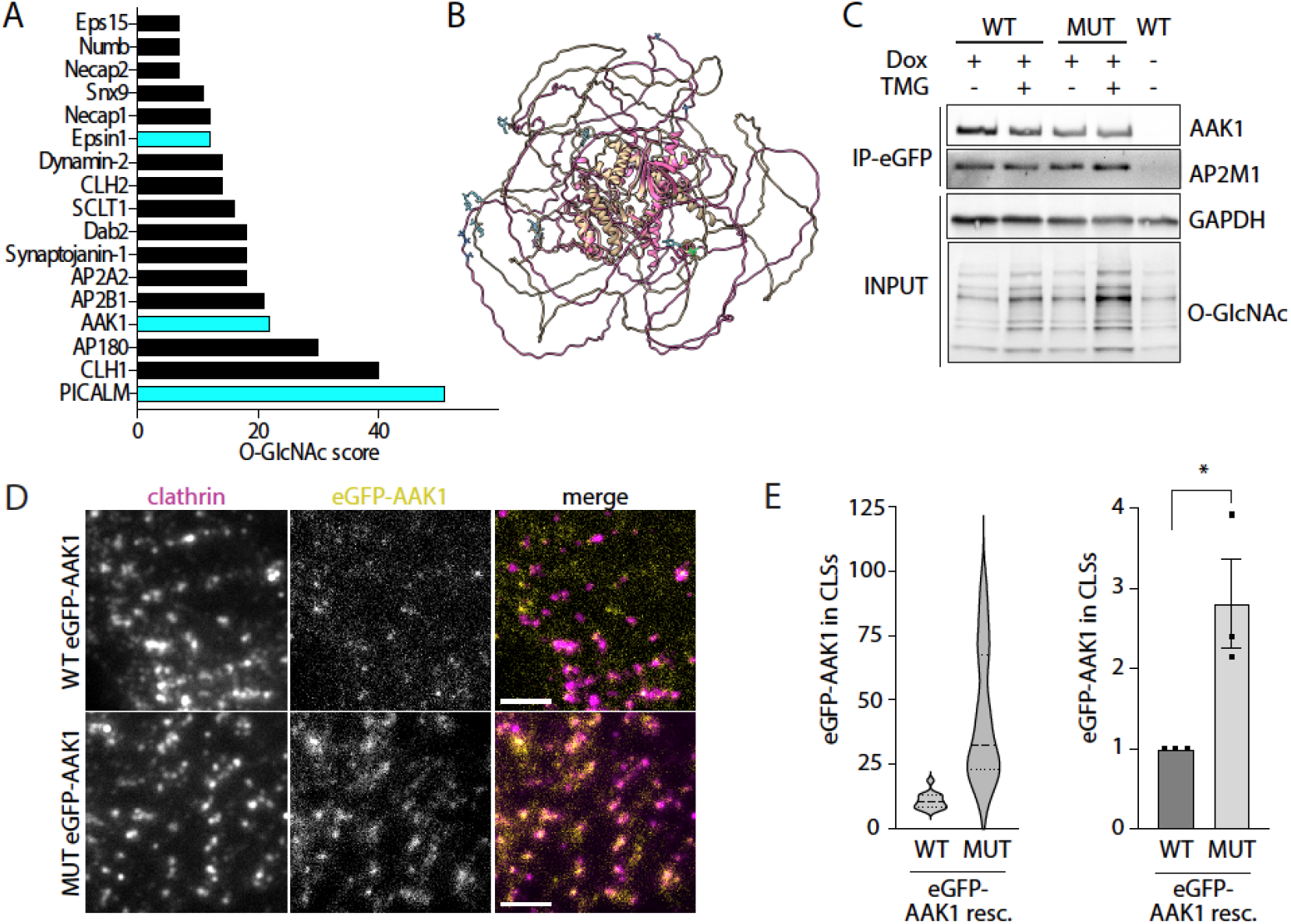
Mutation of predicted O-GlcNAc modification sites alters CCP recruitment of AAK1. (A) Shown is the result of the analysis of (57) showing the probability of O-GlcNAc modification of each endocytic protein (O-GlcNAc score). Highlighted in teal are the three proteins (Epsin1, AAK1, and CALM) examined in further detail here. (B) The structure of AAK1 predicted by AlphaFold, as per *Materials and Methods*. This structure was further decorated with O-GlcNAc at the predicted residues (shown in teal); however, the O-GlcNAc modification was not taken into consideration at the time of structure determination. (C-E) ARPE-19 cells were engineered to stably harbor a transgene for inducible expression of eGFP-tagged AAK1 wild-type (WT) or harboring mutations in the 8 predicted O-GlcNAc modification sites (mut). (C) Following treatment with doxycycline (dox) to induce expression of eGFP-AAK1, samples were subject to immunoprecipitation of eGFP-AAK1 and western blotting as indicated. (D) Cells were treated with siRNA to silence endogenous AAK1. Following siRNA transfection to silence endogenous AAK1 and treatment with dox to induce expression of eGFP-AAK1 proteins, cells were fixed and stained to label clathrin heavy chain, followed by imaging by TIRF and epifluorescence microscopy. Scale 3 μm. (E) Images as in (D) were subjected automated detection and analysis of CLSs as described in *Materials and Methods*. Shown are the measurements of eGFP-AAK1 intensity within CLSs, as the distribution of these values in a representative experiment as a violin plot (left panel), or the mean (bar) ± SE from 3 independent experiments, as well as individual values from each experiment (dots). The total number of CLSs and individual cells (respectively) quantified are as follows: eGFP-AAK1 WT 9645, 33; eGFP-AAK1 mut 9576, 42 (total) *, p < 0.05

Data from the mouse synaptosome revealed AAK1 (Q3UHJ0, AAK1_MOUSE) had 7 O-GlcNAc sites based on a lectin chromatography peptide analysis (59) of which 5 aligned with human AAK1 (Q2M2I8, AAK1_HUMAN). We also identified 3 other O-GlcNAc sites from a new advanced O-GlcNAc database (57). These 8 sites fall in the QP-rich region of AAK1 and not in its C-terminal AP2 interaction domain. In total, we identified 8 serine/threonine residues within human AAK1 that are likely to be modified with O-GlcNAc: T359, T360, T441, S447, T448, T507, S519, S650 (**Figure S2A**). Since experimentally determined structures of AAK1 are currently restricted to its kinase domain, we used AlphaFold2 (60, 61) to predict the structure of full-length AAK1 (**Figure 4B**). The predicted AAK1 structure was consistent with the experimentally determined structures of the AAK1 kinase domain, as alignment of the peptide backbone of the predicted kinases domains and the experimentally determined domains have RMSD values between 0.438 and 0.794 angstroms (**Figure S1)**. This full-length predicted structure revealed that surrounding the highly ordered core of AAK1 that includes the kinase domain, a large portion of the protein is predicted to be comprised of intrinsically disordered regions. Importantly, the residues predicted to be O-GlcNAc modified are found within the intrinsically disordered regions of AAK1 (**Figure 4B**, *O-GlcNAc modification in teal*).

To determine the contribution of specific putative O-GlcNAc-modified AAK1 sites to CCP formation, we developed a knockdown-rescue approach using a mutant of AAK1 (fused to eGFP), which all 8 of these predicted Ser/Thr sites were mutated to Ala (**Figure S2B**). Mutating several potential O-GlcNAc sites in a clustered region is essential as it fits the progressive pattern of OGT modification of its substrates and its reaction mechanism (59). To ensure that the expression of eGFP-AAK1 constructs was controlled to limit possible off-target effects of overexpression, we generated ARPE-19 stable cells using the Sleeping Beauty transposon system (62) that allowed doxycycline-inducible expression at controlled levels. Using this system, we detect wild-type or mutant eGFP-AAK1 expression upon the rescue of AAK1 expression with doxycycline treatment (**Figure S3**).

To assess how the mutations in putative O-GlcNAc modification sites may impact AAK1 function, we first examined its association with AP2, a well-established interactor of AAK1 (18). We immunoprecipitated eGFP-AAK1 wild-type or mutant and probed for the µ2-subunit of AP2 (**Figure 4C**). We detected no differences in the extent of AP2 association with AAK1 mutant comparted to wild type AAK1. This indicates that mutation of putative O-GlcNAcylation sites on AAK1 does not grossly disrupt its structure or association with AP2, consistent with a computational analysis of the effect of the mutations showing minimal impact on AAK1 secondary structure (**Figure S4**).

To elucidate how the putative O-GlcNAc modification sites impact AAK1 recruitment to clathrin structures, we performed TIRF microscopy on cells treated with siRNA to silence endogenous AAK1, followed by induction of eGFP-AAK1 proteins, immunostaining to label clathrin, and imaging by TIRF microscopy (**Figure 4D**). We examined these images with automated detection and analysis of CLSs, and found that mutant eGFP-AAK1 was recruited to clathrin structures much more robustly than wild-type eGFP-AAK1, both when examining the distribution of individual CLSs in a single representative experiment (**Figure 4E**, *left panel*) and in multiple independent experiments (**Figure 4E**, *right panel*). While there was no significant difference between eGFP-AAK1 wild type and mutant expressing cells with respect to clathrin intensity per CLS (**Figure S5A**), we observed a robust increase in clathrin intensity per CLS upon the slight overexpression of eGFP-AAK1 wild type achieved by this knockdown-rescue strategy (**Figure S5B**). This is consistent with an increase in AAK1 expression and thus increased AAK1 recruitment to CCPs leading to an increase in CCP size and suggests that there is a limit to the ability of AAK1 to increase CCP size.

Collectively, these results indicate that the level of AAK1 recruited to clathrin structures may control the size of CCPs. Furthermore, our results show that the mutation of the eight putative O-GlcNAc modification residues robustly increases AAK1 to clathrin structures, suggesting that OGT and O-GlcNAcylation of AAK1 may suppress AAK1 recruitment to CCP. We attempted to detect O-GlcNAc modification AAK1 by antibody based-detection of O-GlcNAc residues but were unable to resolve O-GlcNAcylation of AAK1 by this method, perhaps due to this approach having some sequence-specificity (63). Nonetheless, the previous determination of AAK1 O-GlcNAc modification (51) and the functional effects on AAK1 recruitment to clathrin structures upon mutation of putative O-GlcNAc modified residues (**Figure 4**) suggest that AAK1 may be subject to O-GlcNAc modification to suppress AAK1 recruitment to CCPs.

In addition to AAK1, CALM and Epsin were also identified as proteins with substantial composition of intrinsically-disordered regions and a high likelihood of O-GlcNAc modification (**Figure 4A**). We next examined how OGT silencing can impact the recruitment of CALM or Epsin to clathrin structures, using staining of each protein in cells expressing eGFP-clathrin, followed by TIRF microscopy (**Figure 5A-B**) and CLS analysis. We previously validated Epsin antibodies (64) and validated CALM antibodies (**Figure S6**) for use in TIRF microscopy. The level of CALM recruited to CLSs was significantly increased upon OGT silencing when examining individual CLS measurements in a representative experiment (**Figure 5C**) and in multiple independent experiments (**Figure 5D**). Since OGT silencing increases the recruitment of eGFP-clathrin in clathrin structures, we asked whether if the increase in CALM recruitment to CLSs scales with clathrin. The ratio of CALM to eGFP-clathrin intensity in CLSs was unchanged upon OGT silencing when examining individual CLS measurements in a representative experiment (**Figure 5E**) and in multiple independent experiments (**Figure 5F**). This suggests that while OGT silencing triggers an increase in CALM recruitment to CLSs, this increase in CALM may reflect an increase size of these structures and not a specific effect on CALM recruitment *per se*.

**Figure 5.**
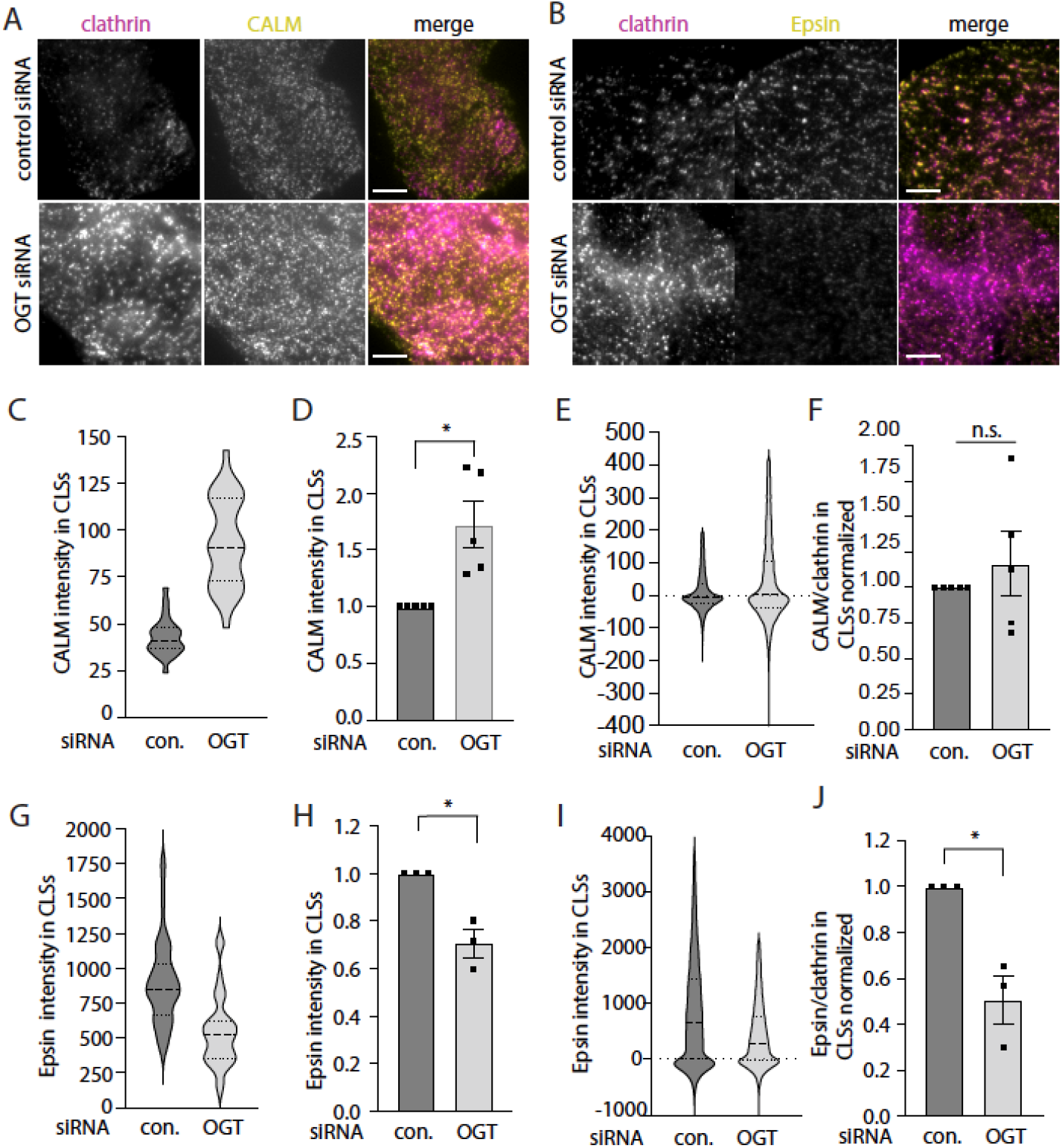
OGT silencing distinctly impacts recruitment of CALM and Epsin to CCPs. ARPE-19 cells stably expressing eGFP-clathrin light chain were treated with siRNA to silence OGT or non-targeting (control) siRNA, subjected to immunofluorescence staining to detect CALM or Epsin1, and then imaged using TIRF microscopy. Shown are representative TIRF images for CALM (B) or Epsin1 (C) experiments. scale 5 µm. (C-J) Images as in (A-B) were subjected automated detection and analysis of CLSs as described in *Materials and Methods*. (C-D) The measurements of CALM intensity within CLSs, shown as the distribution of these values in a representative experiment as a violin plot (C), or the mean (bar) ± SE from 5 independent experiments, as well as individual values from each experiment (dots) (D). (E-F) The ratio of CALM/eGFP-clathrin intensities within CLSs, shown as the distribution of these values in a representative experiment as a violin plot (E), or the mean (bar) ± SE from 5 independent experiments, as well as individual values from each experiment (dots) (F). (G-H) The measurements of Epsin1 intensity within CLSs, shown as the distribution of these values in a representative experiment as a violin plot (G), or the mean (bar) ± SE from 5 independent experiments, as well as individual values from each experiment (dots) (H). (I-J) The ratio of Epsin1/eGFP-clathrin intensities within CLSs, shown as the distribution of these values in a representative experiment as a violin plot (I), or the mean (bar) ± SE from 5 independent experiments, as well as individual values from each experiment (dots) (J). The total number of CLSs and individual cells (respectively) quantified are as follows, for CALM staining: control siRNA 95762, 134; OGT siRNA 106993, 139; for Epsin1 staining: control siRNA 61718, 85; OGT siRNA 55792, 79 *, p < 0.05

In contrast to the effect of OGT silencing on CALM recruitment, loss of OGT triggered a loss of Epsin from CLSs, when examining individual CLSs measurements in a representative experiment (**Figure 5G**) and in multiple independent experiments (**Figure 5H**). This loss of Epsin recruitment to CLSs was retained when examining the ratio of Epsin to clathrin (**Figure 5I-J**). This indicates that OGT does not act similarly on all endocytic accessory proteins; instead, while OGT and protein O-GlcNAcylation suppress the recruitment of clathrin, AAK1, and CALM to clathrin structures, OGT and protein O-GlcNAcylation enhance Epsin recruitment to clathrin structures. The selective control of recruitment of different endocytic accessory proteins by OGT is consistent with cargo-selective effects on the recruitment of EGFR and TfR to clathrin structures (**Figure 3**).

Our results indicate that loss of OGT and cellular O-GlcNAc protein modification suppresses eGFP-AAK1 recruitment to clathrin structures and leads to an increase in CCP size and loss of Epsin recruitment therein. To determine if AAK1 may be functionally required for the alterations in CCP size or Epsin recruitment seen upon perturbation of OGT, we first examined the effect of inhibition of AAK1 by treatment with the small molecule inhibitor LP-935509. As expected, treatment with LP-935509 led to a loss of phosphorylation of the µ2 subunit of AP2 (**Figure S7**), establishing effective inhibition of AAK1 kinase activity.

To determine if AAK1 contributes to the increase in CCP size upon loss of OGT, we examined the effect of LP-935509 treatment following OGT silencing in cells expressing eGFP-clathrin and imaged by TIRF (**Figure 6A**) and widefield epifluorescence (**Figure 6B**) microscopy. Consistent with our results so far, automated detection and analysis of CLSs in TIRF microscopy images showed that OGT silencing triggered an increase in eGFP-clathrin intensity in CLSs, and this was not affected in cells treated with LP-935509 (**Figure 6C**). Since the intensity of eGFP-clathrin within CLSs in TIRF images can be impacted by both the amount of eGFP-clathrin molecules recruited per structure or the curvature of CLSs, we also examined the intensity of CLSs in widefield epifluorescence microscopy images, which are only impacted by the amount of eGFP-clathrin per CLS and not their curvature. While OGT silencing increased eGFP-clathrin intensity in CLSs in epifluorescence images, this effect was ablated by treatment with LP-935509 (**Figure 6D**). That the intensity of eGFP-clathrin in CLSs in TIRF images remains increased upon treatment with LP-935509, while this value is suppressed in widefield epifluorescence images suggests that LP-935509 treatment impairs both the increase in size of CLSs upon OGT silencing and the acquisition of curvature of these structures. Taken together, these results indicate that AAK1 is required for the increase in clathrin structure size upon loss of OGT, suggesting that the increased recruitment of AAK1 to clathrin structures may be functionally required for the alterations in clathrin structure assembly seen upon loss of OGT.

**Figure 6.**
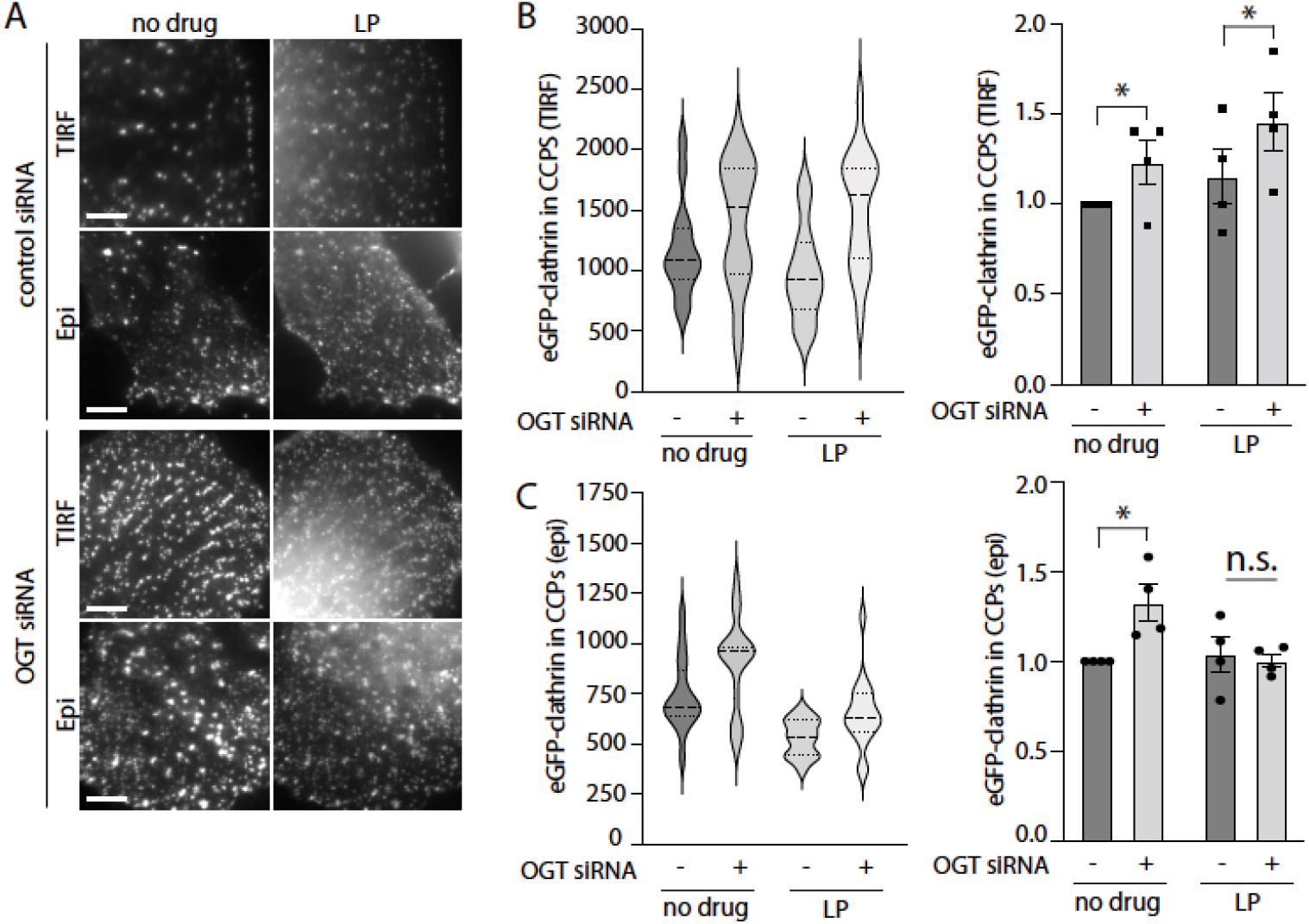
Inhibition of AAK1 with LP-935509 reverses the effect of OGT silencing on CCP size. ARPE-19 cells stably expressing eGFP-clathrin light chain were treated with siRNA to silence OGT or non-targeting (control) siRNA. Then, immediately prior to fixation, cells were treated with 5 μM LP-935509 (LP) or vehicle control (no drug) for 3 hours, and then imaged using TIRF and epifluorescence microscopy. Shown are representative images obtained by TIRF (A) or epifluorescence (B) microscopy. scale 5 µm. Images as in (A-B) were subjected automated detection and analysis of CLSs as described in *Materials and Methods*. Shown is the intensity of eGFP-clathrin within CLSs detected in TIRF (C) or epifluorescence (D) images, in each case showing the distribution of values within a single representative experiment as a violin plot (*left panels*) and the mean (bar) ± SE from 5 independent experiments, as well as individual values from each experiment (dots) (*right panels*). The total number of CLSs and individual cells quantified (respectively) are as follows, control siRNA, no drug 6335, 46; control siRNA, LP-935509, 9291, 76; OGT siRNA, no drug 9309, 70; OGT siRNA, LP-935509, 10063, 76 ; *, p < 0.05

### Nutrient availability impacts O-GlcNAc protein modification and clathrin structure formation

The impact of OGT and cellular protein O-GlcNAc modification suggest that signals that regulate O-GlcNAc modification may exert control over CCP formation. Cellular protein O-GlcNAcylation is highly sensitive to metabolic flux through the hexosamine biosynthetic pathway, and thus the availability of nutrients, in particular glucose and glutamine (65). We next aimed to resolve if the availability of glucose and glutamine may regulate CCP formation. To do so, we incubated cells for 4h in either media devoid of glucose and glutamine (which we term “low GG”) or the same base media supplemented with 10 mM glucose and 2.5 mM glutamine (which we term “high GG”). Comparing the levels of cellular O-GlcNAc modification shows that these conditions substantially impacted the total levels of cellular protein O-GlcNAc modification (**Figure 7A**).

**Figure 7.**
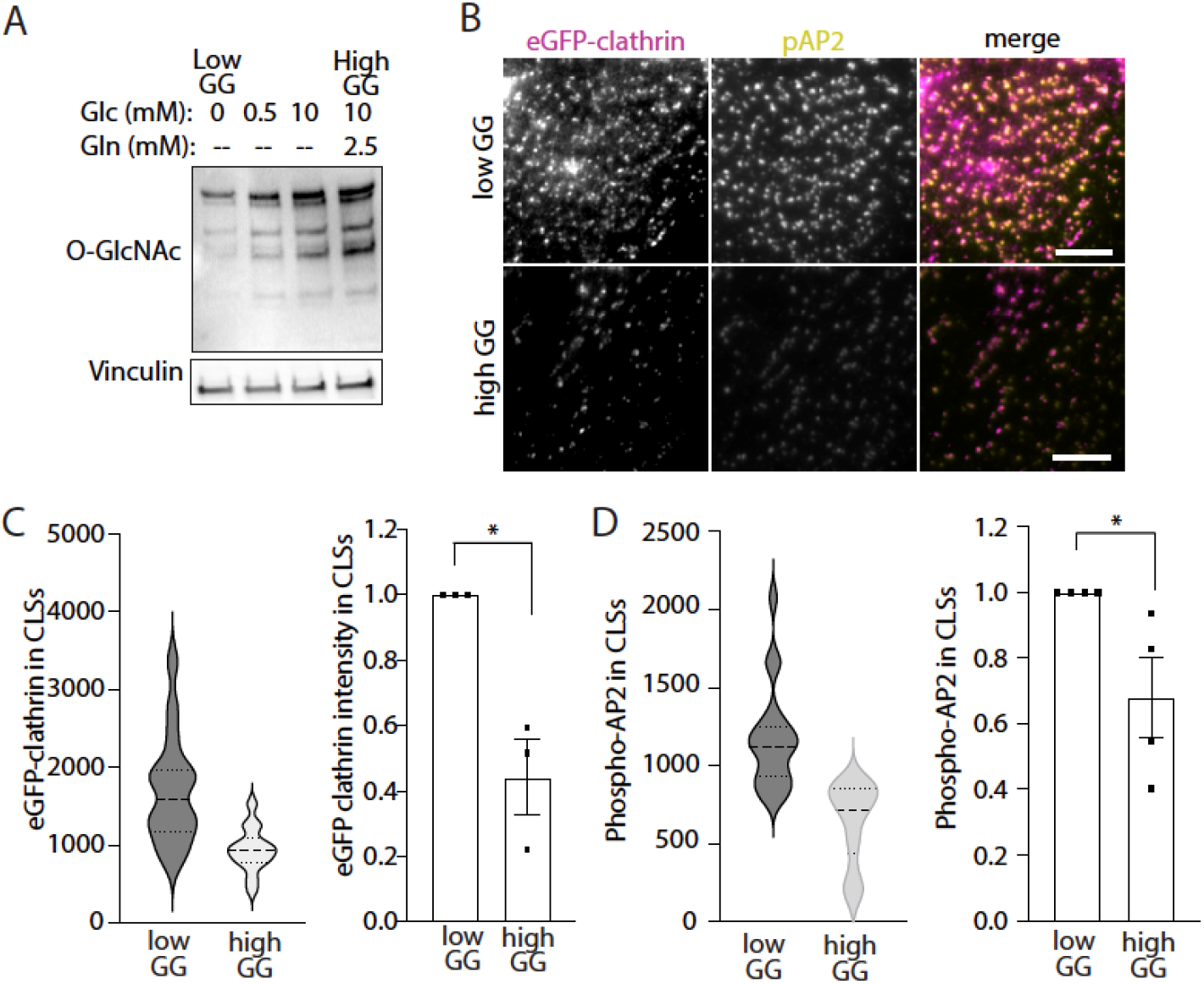
Nutrient availability impacts O-GlcNAcylation and CCP size. ARPE-19 cells stably expressing eGFP-clathrin were incubated in media containing glucose and glutamine as indicated for 4 h (low GG: 0 mM glucose and 0 mM glutamine; high GG: 10 mM glucose and 2.5 mM glutamine). (A) Immunoblots of whole cell lysates probed with antibodies as indicated. (B) Following lowGG/high GG treatment for 4h, cells were fixed, subjected to staining to detect phosphorylated AP2, and then imaged by TIRF microscopy. Scale 5 µm. (C-D) Images as in (B) were subjected automated detection and analysis of CLSs as described in *Materials and Methods*. Shown is the intensity of eGFP-clathrin (C) or pAP2 (D) within CLSs detected in TIRF microscopy images, in each case showing the distribution of values within a single representative experiment as a violin plot (left panels) and the mean (bar) ± SE from 3 independent experiments, as well as individual values from each experiment (dots) (right panels). The total number of CLSs and individual cells quantified (respectively) are as follows, low GG 45763, 69; high GG 45544, 66; *, p < 0.05

To determine how nutrient availability impacts CCP formation and AAK1 activity, we incubated cells stably expressing eGFP-clathrin for 4h in either low-GG or high-GG media. We then stained cells for phospho-AP2 (the substrate of AAK1), followed by imaging by TIRF microscopy (**Figure 7B**). We then examined images through automated detection and analysis of CLSs. Consistent with the effect of OGT silencing that increased CCP size, we observed that low-GG conditions that exhibit reduced protein O-GlcNAcylation had a significantly elevated intensity of eGFP-clathrin in CLSs, as observed in CLSs measurements in a single representative experiment (**Figure 7C,** *left panel*), as well as in multiple independent experiments (**Figure 7C,** *right panel*). We observed similar results for phospho-AP2 (**Figure 7D**). Together, these results indicate that the availability of glucose and glutamine exert significant effects on CCP formation that are correlated with the regulation of CCP formation by protein O-GlcNAcylation.

## Discussion

Understanding how cells adapt their endocytic membrane traffic to their environment under different metabolic conditions is central to understanding the contribution of membrane traffic under distinct physiological contexts. We investigated how the nutrient-responsive O-GlcNAc modification modulates clathrin-mediated endocytosis. We identified that O-GlcNAc affects the endocytosis machinery of clathrin-mediated endocytosis using multiple independent approaches, including siRNA silencing of OGT, pharmacological manipulation of OGA, and use of a mutant of AAK1 that is predicted to lose O-GlcNAc modification. To our knowledge, we provide the first direct functional evidence that clathrin-mediated endocytosis spatiotemporal dynamics are modulated by an intrinsic metabolic signal: protein O-GlcNAcylation. This provides further evidence as to the versatility and adaptability of endocytic traffic via O-GlcNAc modification.

The post-translational modification with O-GlcNAc has been difficult to detect and quantify on a per protein basis, as it is highly dynamic, present at sub-stoichiometric amounts on nucleocytoplasmic proteins, and is labile due to its chemical structure (55, 56). The current methods to detect and quantify O-GlcNAc modifications on proteins come with many limitations and it is difficult to identify specific sites of O-GlcNAc modification (56). We were not able to detect O-GlcNAc modification of AAK1 following immunoprecipitation either by mass spectrometry or western blotting. However, not being able to detect O-GlcNAc modification is not uncommon for proteins modified with O-GlcNAc. Recently, a similar mutagenesis strategy was used (66) for creating mutants of galectin 3 that had alteration of 4 Ser/Thr residues predicted by *in silico* analysis to be O-GlcNAc modified. These sites in galectin 3 were similarly mutated to Ala without clearly identifying the sites of O-GlcNAc modification. Thus, the functional outcome of mutations of candidate O-GlcNAc sites is compelling even in the absence of evidence of O-GlcNAc modification (52).

AAK1 was previously shown to be subject to O-GlcNAc modification (51, 59). While the specific residues that are subject to O-GlcNAc modification are not known, we found that a mutant of AAK1 with eight predicted O-GlcNAc modification Ser/Thr residues changed to Ala exhibited significant alterations in recruitment to CCPs (**Figure 4**). Specifically, the mutant AAK1 exhibited robustly increased recruitment to CCPs despite expression at similar levels as wild-type AAK1 (**Figure S3**). Based on the findings in cells expressing eGFP-AAK1, loss of O-GlcNAc modification in the mutant AAK1 leads to enhanced recruitment of AAK1 to CCPs, which would be expected to enhance phosphorylation of AP2 and promote CCP assembly and stabilization (5, 21). Hence, the loss of the putative O-GlcNAc modification sites on AAK1 increases AAK1 localization to clathrin structures, which is consistent with the enhanced initiation rate and size of CCPs that we observed upon silencing OGT.

Notably, even the mild overexpression of the wild-type eGFP-AAK1 increased AP2 phosphorylation and CCP size, which was not further enhanced by expression of the eGFP-AAK1 with mutations in the putative O-GlcNAcylation sites. Hence, these results indicate that increased recruitment of AAK1 to CCPs does increase AP2 phosphorylation and CCP size, but that there is a limit to the extent of this increase that is already reached by the levels of AAK1 recruited to CCPs in the wild-type eGFP-AAK1 expressing cells. Taken together, these results suggest a model in which the loss of O-GlcNAc modification of AAK1, either by mutation of putative O-GlcNAcylation sites on AAK1 or by silencing OGT, may promote recruitment of AAK1 to CCPs, which in turn may promote AP2 phosphorylation and enhance CCP assembly and increase CCP size.

The putative O-GlcNAc modification sites are in regions of AAK1 that are predicted to be intrinsically disordered. The mutagenesis of eight Ser/Thr sites to Ala did not affect the predicted disordered regions of AAK1, indicating that the structure of AAK1 was not grossly impacted in this mutant, which is corroborated by the interaction with AP2 being retained in the AAK1 mutant (**Figure 4C**). It is not clear how O-GlcNAc modification of AAK1 may regulate the recruitment of this kinase to CCPs. It may be tempting to speculate that the intrinsically disordered regions of AAK1 could contribute to liquid-liquid phase separation within CCPs, and that O-GlcNAcylation of AAK1 may thus limit AAK1 partitioning within such a structure. It is also possible that O-GlcNAcylation of AAK1 may alter secondary structure or otherwise alter protein-protein interactions of AAK1 with clathrin resident proteins other than AP2.

While our results suggest that O-GlcNAc modification of AAK1 may regulate the recruitment of this kinase to CCPs and thus the properties of CCPs, it is also possible that other CCP resident proteins are modified with O-GlcNAc, impacting CCP properties as seen in experiments involving perturbations of OGT or OGA. It is also possible that O-GlcNAc-based regulation of CCP dynamics may be indirect, with broad alterations in proteins distal to CCPs impacting endocytic assemblies by other mechanisms such as changes in gene expression. While a systematic analysis of all CCP proteins to resolve other possible O-GlcNAc modifications is beyond the scope of this study, future research that can further resolve how O-GlcNAc post-translational modification may regulate CME and endocytic membrane traffic will be very informative.

Our results indicate that CCP initiation, assembly and stabilization, as well as cargo recruitment, is robustly controlled by protein O-GlcNAcylation. Interestingly, another study found that the clathrin-dependent internalization was unaffected by silencing of OGT or OGA (66). In contrast, we found that suppression of OGT and OGA substantially impacted CCP initiation and assembly. Specifically, suppression of OGT enhanced CCP initiation and size, which would contribute to enhanced endocytosis. At the same time, OGT silencing also lead to decreased CCP lifetimes that may reflect enhanced abortive turnover of CCPs without internalization, and a reduction of TfR recruitment to CCPs, both of which should suppress endocytosis. Hence, it is possible that OGT and OGA perturbations impact CCP dynamics in a manner to limit changes to the internalization of some receptors under a range of metabolic conditions. Alternatively, or in addition, the integration of effects controlled by O-GlcNAc modification may have limited effects on the internalization of some cargo (e.g. TfR), while significantly impacting the internalization of others. Consistent with this, inhibition of AAK1 leads to modest (∼25%) inhibition of TfR internalization (5, 21), while AAK1 may have much more significant control over that of LRP6 (67).

Our observation that the recruitment of EGFR to CCPs upon OGT silencing scales with the amount of clathrin, while that of TfR does not, suggests that O-GlcNAc modification may selectively impact the internalization of EGFR and TfR. This possible difference in regulation of CME of TfR and EGFR by protein O-GlcNAcylation could occur as a result of direct and distinct O-GlcNAc modification of each receptor, as has been reported for EGFR (68). Alternatively, as TfR and EGFR reside largely in distinct CCPs (15), EGFR and TfR internalization may be differently regulated by O-GlcNAc modification as a result of distinct outcomes on subsets of CCPs. Interestingly, O-GlcNAc modification of hepatocyte growth factor regulated tyrosine kinase substrate (HGS) alters EGFR intracellular traffic by suppressing ligand-stimulated EGFR degradation, leading to sustained signaling (69). As CCPs at the plasma membrane are also important regulators of EGFR signaling (15, 53, 54, 64,70,71), protein modification by O-GlcNAc may act to regulate EGFR membrane traffic and signaling at multiple levels.

Resolving how the robust modulation of CCP initiation, size and lifetimes by O-GlcNAcylation impacts the internalization and membrane traffic of the diverse proteins at the cell surface will require systematic methods to study cell surface protein abundance. Such approaches have been done to study the impact of AP2 perturbation (72), or AMPK activation (39). Using similar approaches to study the effect of OGT or OGA perturbation may allow identification of a cohort of cell surface proteins that exhibit internalization with preferential regulation by protein O-GlcNAcylation.

In conclusion, we find that protein O-GlcNAc modification exerts control over the formation, assembly and maturation of CCPs. As such, O-GlcNAc modification may be an important mediator of the control of endocytic membrane traffic by cellular metabolic state (13, 14). Endocytic membrane traffic controls the access of myriad cell surface proteins to their substrates in the extracellular milieu. Thus, further resolving of metabolic signals rely on protein O-GlcNAc modification and other mechanisms to control membrane traffic and this cell surface proteins will continue to be an important area of investigation.

### Experimental Procedures

#### Materials

Details about antibodies used for western blotting and immunofluorescence experiments are in **Table S1**. Fluorophore-conjugated or horseradish peroxidase (HRP) secondary antibodies were from Jackson ImmunoResearch (West Grove, PA). Rhodamine-EGF was generated in-house as previously described by (73).

#### Cell Culture

Wild-type human retinal pigment epithelial (ARPE-19) cells were cultured in Dulbecco’s Modified Eagle’s Medium/F-12 (DMEM/F12; Thermo Fisher Scientific, Mississauga, ON) supplemented with 10% fetal bovine serum (Thermo Fisher Scientific), and an antibiotic cocktail consisting of 100 U/mL penicillin and 100 μg/mg streptomycin (Thermo Fisher Scientific). Cells were incubated at 37°C and 5% CO_2_ until they reached 80% confluency and were suitable for passaging. Cells were passaged every 2-3 days, washed with sterile phosphate-buffered saline (Sigma Aldrich, Oakville, ON) and lifted with 0.25% Trypsin-EDTA (Thermo Fisher Scientific).

#### Stable transfections using Sleeping Beauty transposon system

pSB-tet-BP was a gift from Eric Kowarz (Addgene plasmid # 60496 ;http://n2t.net/addgene:60496; RRID:Addgene_60496) (62). pCMV(CAT)T7-SB100 was a gift from Zsuzsanna Izsvak (Addgene plasmid # 34879 ; http://n2t.net/addgene:34879; RRID:Addgene_34879) (74). An oligonucleotide encoding eGFP fused to AAK1 were generated by BioBasic Inc (Markham, ON, Canada), using the sequence of eGFP, followed by the sequence encoding a spacer peptide (GGG GGG TCT GGT GGC AGT GGA GGG GGA TCC), followed by the sequence of human AAK1, as per accession number NM_Q2M2I8. This oligonucleotide sequence was subcloned into pSB-tet-BP to generate pSB-tet-BP-eGFP-AAK1(WT). From this plasmid, mutant AAK1 (S447, T507, S519, T359, T360, Thr448, Thr441, Ser650 to Ala) constructs were derived by site-directed mutagenesis by BioBasic Inc (**Figure S2B**). pSB-tet-BP plasmids encoding various AAK1 WT and AAK1 mutant constructs alongside pCMV(CAT)T7-SB100 were co-transfected into ARPE-19 cells using FuGene HD transfection reagent, as per manufacturer’s protocol (Promega, Madison, WI), followed by selection of stably engineered cells in media supplemented with 2 µg/mL puromycin for a period of 2-3 weeks.

pSBtet-BP stable cells were kept in cell culture media as indicated above but were incubated with 10% fetal bovine serum without tetracycline and maintained in 2 µg/mL puromycin. For the induction of AAK1-GFP WT or mutant Doxycycline (Dox) was used at a final concentration of 1µM in the cell culture media for 24 hours before use of cells for different downstream applications.

#### Western blotting

Western blotting was performed as previously described (75). After incubation and pharmacological treatments as indicated, cells were rapidly cooled by washing with ice-cold PBS and then lysed in 2X-Laemelli Sample buffer (0.5M Tris pH 6.8, Glycerol, 10% SDS) supplemented with 1 mM sodium orthovanadate, 10 mM okadaic acid and 20 mM of protease inhibitor. Whole-cell lysates were then syringed 5 times with a 27.5-gauge syringe. Lysates were then heated at 65°C for 15 minutes under reducing conditions (with supplementation of lysis buffer with 10% beta-mercaptoethanol and 5% bromophenol blue), resolved by SDS-PAGE, and transferred to 0.2 µm pore PVDF membrane (Pall Life Science, Port Washington). The membrane was blocked with 3% BSA and then incubated with indicated antibodies in 1% BSA at 1:1000 dilution at 4°C overnight. The membrane was then subjected to a secondary HRP-conjugated antibody (with 1% BSA) and left to shake for 1 hour at RT before imaging. Images were obtained using a BioRad ChemiDoc Touch Imaging System upon soaking membranes in Luminata Crescendo HRP substrate (Millipore Sigma).

#### siRNA transfections

ARPE-19 cells were subject to siRNA gene silencing for a specific target (OGT, CALM, AAK1) or treated with a non-targeting (control) siRNA as previously described (53, 75–77). Custom siRNAs were synthesized to target specific transcripts with sense strand sequences (with overhang sequence in parentheses) as follows: OGT: GAA GAA AGU UCG UGG CAA A (UU), CALM: CCU CAU ACC UCU UUA ACA A (UU), AAK1: GGG AAA GUC AGG UGG CAA U (UU), control: CGU ACU GCU UGC GAU ACG G (UU). siRNA gene silencing was performed using Lipofectamine RNAiMAX (Life Technologies, Carlsbad, CA), as per the manufacturer’s instructions. Each siRNA construct was transfected at 220 pmol/l precomplexed and incubated in the transfection reagent in Opti-MEM (Gibco) for 4 h. After the 4 h incubation, cells were washed and replaced in a regular 10% FBS DMEM/F12 growth medium. siRNA transfections were performed twice (72 h and 48 h prior to experiments) except for OGT, which was performed only one time, 48h before experiments.

#### Purification of GFP-fusion proteins

*Escherichia coli* BL21 cells transformed with GFP-binding V_H_H domain, GFP-binding protein (GBP) in pHEN6 vector were a gift from Dr. Greg Fairn (Dalhousie University). The cells were induced with 1mM isopropyl-β-D-1-thiogalactopyranoside for 3-4 hours in 30°C. Bacterial cells were harvested by centrifugation (10 min at 3000g) and the pellet was resuspended in lysis buffer (50 mM Tris pH 7.5, 300 mM NaCl, 10 mM imidazole, 5% (v/v) Glycerol). The cell suspension was sonicated at 4°C for 2 minutes with 20 s on, and 59.9 s off, with 15% amplitude. After centrifugation (20 min at 14,000 rpm) the soluble protein was loaded onto preequilibrated 1 mL NiNTA resin and purified. The His-tagged GBP was eluted by using NiNTA elution buffer (20 mM Tris pH 7.5, 300 mM NaCl, 250 mM Imidazole, 5% (v/v) Glycerol). Elution fractions containing GBP were pooled and dialyzed with Slide-A-Lyzer Dialysis Cassette G2 (3,500 MWCO, 3 mL capacity) into Dulbecco’s Phosphate Buffered Saline (D8537) to remove the imidazole. The protein concentration was measured with a BCA.

The purified GBP was coupled to N-hydroxysuccinimide activated Sepharose 4 Fast Flow (GE Healthcare) as described previously by (78) with certain modifications. The coupling solution/medium were incubated at 0.5:1 while rotating for 3 hours at room temperature. After coupling, the non-reacted groups on the medium were blocked by 0.1 M Tris-HCL, pH 8.5 for 2-3 hours. The washes were alternatively between two different buffers (high and low pH) 0.1 M Tris-buffered saline pH 8.0 and 0.1 M acetate 500 mM NaCl pH 4.5. The final affinity medium, referred to as GFP-NanoTrap was stored in TBS with 20% ethanol.

AAK1-GFP WT or AAK1-GFP mutant or eGFP-clathrin cells were seeded onto 10 cm dishes until 80-85% confluent, then homogenized in 400 µL cold lysis buffer (20 mM Tris/HCl, pH 7.5, 150 mM NaCl, 0.5 mM EDTA, 2 mM PMSF, 0.5% Nonidet P-40). After a centrifugation step (10 min at 14,000 rpm at 4°C) the supernatant was adjusted with dilution buffer (20 mM Tris/HCl, pH 7.5, 150 mM NaCl, 0.5 mM EDTA, 2 mM PMSF) and incubated in 100 µL of GFP-NanoTrap. The GFP-NanoTrap and protein lysate was incubated overnight at 4°C while rotating. The next day, the samples were washed with dilution buffer and then with wash buffer 2 (20 mM Tris/HCl, pH 7.5, 300 mM NaCl, 0.5 mM EDTA, 2 mM PMSF) just before elution. The elution of GFP-protein was done by adding LSB directly to the cell lysate and incubating it at 65°C for 5 minutes. After this, the samples were resolved by western blotting.

#### Experiment seeding and pharmacological treatments

ARPE-19 cells stably expressing eGFP-clathrin light chain were seeded onto glass coverslips and in some cases subjected to siRNA transfection as indicted (**Figures 1-2, 3, 5-7**). For experiments presented in **Figure 2**, Cells were incubated for 4h at 20 μM Thiamet G for or in vehicle (dH_2_0) prior to live-cell microscopy. For experiments presented in **Figure 3**, following 1 h serum deprivation, cells were treated with 20 ng/ml rhodamine EGF and 10 μg/ml A657-Tfn for 5 min at 37C, followed by immediate fixation in 4% PFA. For AAK1 knockdown-rescue experiments (**Figure 4D-E**), pSBTet-AAK1-GFP WT and pSBTet-AAK1-GFP mutant cells were seeded onto glass coverslips the day before siRNA transfection. Then, after two rounds of siRNA transfection to AAK1, doxycycline was added to the cells for the final 24h before the experiment. For experiments presented in **Figure 6**, cells were either incubated for 3h at 5μM of LP-935509 or in vehicle control (0.1% (v/v) DMSO) prior to the fixation. For experiments presented in **Figure 7**, cells were seeded onto 6-well plates for 24 prior to incubation in media without glucose, glutamine and no phenol red (catalog no. A1443001) supplemented with 10% dialyzed fetal bovine serum (Thermo Fisher Scientific), and glucose and/or glutamine, as indicated, for 4h prior to fixation.

#### Immunofluorescence staining

For detection of total cellular protein (all immunofluorescence experiments), after treatments as indicated cells were fixed 4% paraformaldehyde for 30 min, followed by quenching of fixative in 100 mM glycine, cell permeabilization in 0.1% Triton X-100 (all solutions made in PBS), and then blocking in 3% BSA (Thermo Fisher Scientific). Subsequently, cells were stained with primary and secondary antibodies as indicated and retained within an aqueous medium for imaging by TIRF microscopy.

#### Microscopy and image analysis

##### Image acquisition

Microscopy was performed using a Quorum (Guelph, ON, Canada) Diskovery microscope that is comprised of a Leica DMi8 microscope equipped with a 63×/1.49 NA TIRF objective with a 1.8× camera relay (total magnification 108×). Imaging was done using 488-, 561, and 637-nm laser illumination and 527/30, 630/75, and 700/75 emission filters. The microscope was operated in total internal reflection fluorescence microscopy (TIRF-M) or widefield epifluorescence microscopy modes, as indicated. Images were acquired using a Zyla 4.2Plus sCMOS camera (Oxford Instruments). Fixed-cell TIRFM imaging was done at room temperature with samples mounted in PBS. For live-cell imaging experiments (**Figures 1,2**), cells were maintained at constant 37C during imaging, in phenol-free DMEM/F12 media (Gibco) supplemented with 20 mM HEPES.

##### Detection and analysis of CCP dynamics in time-lapse image series

Automated detection, tracking and analysis of CCPs (as in **Figures 1-2**) was as previously described (4–6, 64) following time-lapse TIRF microscopy of RPE cells stably expressing eGFP-clathrin light chain. Diffraction-limited clathrin structures were detected using a Gaussian-based model method to approximate the point-spread function (6), and trajectories were determined from clathrin structure detections using u-track (79). sCLSs were distinguished from *bona fide* CCPs based on unbiased analysis of clathrin intensities in the early CCP stages (5, 6). Both sCLSs and CCPs represent nucleation events, but only *bona fide* CCPs represent structures that undergo stabilization, maturation and in some cases scission to produce vesicles (5, 6). We report the sCLS nucleation, CCP initiation, CCP lifetime distribution, and the density of persistent CCPs, as well as the intensity of eGFP-CLC within structures as the ‘plateau intensity’ of eGFP-clathrin within these structures. CCPs exhibit several phases including initiation, growth/assembly, plateau and disassembly/scission (80). Here we define the ‘plateau intensity’ of eGFP-CLC in TIRF or epifluorescence microscopy images as the mean fluorescence of eGFP-clathrin within each detected clathrin structure, measured within timepoints corresponding to 30% and 70% of the total lifetime of that structure, during which time CCPs exhibit minimal growth or disassembly (64). Because CCPs are diffraction-limited objects, the amplitude of the Gaussian model of the fluorescence intensity of eGFP-CLC informs about CCP size (**Figures 1E,F; 2E,F**). All measurements were subjected to ANOVA followed by Tukey post-test with a threshold of p < 0.05 for statistically significant differences between conditions.

##### Detection and analysis of CLS in fixed cell samples

CLSs were detected and quantified using custom software developed in Matlab (MathWorks, Natick, MA), as previously described (6, 15, 64). Briefly, diffraction-limited clathrin structures were detected using a Gaussian-based model method to approximate the point-spread function of eGFP-CLCa, or antibody-labeled clathrin in TIRF-M images. The fluorescence intensity corresponding to the secondary channel such as that of a second label (e.g. CALM, EPSIN, AAK1) or of clathrin within epifluorescence images within CLSs was determined by the amplitude of the Gaussian model for the appropriate fluorescent channel for each structure. As such, the measurements of protein intensity within CLSs represent enrichment of the corresponding signal relative to the local background fluorescence in the vicinity of the detected CLS. Measurements (mean levels of various proteins within specified CLS subset for each cell) were subjected to either two-sided student’s t-test or ANOVA followed by Tukey post-test, with a threshold of p < 0.05 for statistically significant differences between conditions.

#### Structure analysis of AAK1

AlphaFold 2 (60, 61) was used via collabfold (Mirdita et al., 2022) to predict the three-dimensional structure of AAK1 (961 amino acids) using the sequence NM_Q2M218 and an 8 alanine (S447, T507, S519, T359, T360, T448, T441, S650) mutant sequence (mutant AAK1). Structures were visualized using ChimeraX (Pettersen et al., 2021; Goddard et al., 2018) and O-GlcNac residues were added by overlaying sugar onto S/T hydroxyl groups. Structural predictions of the low complexity regions were independently identified using Protein Disorder Prediction System Server (PrDOS) (Ishida and Kinoshita, 2007). A scatter plot was generated using the disorder predictions of both the WT and the mutant forms of AAK1 (**Figure S4**). Experimentally determined crystal structures available to date of the kinase domain from human AAK1 were modelled to the predicted structure of AAK1 using ChimeraX (Pettersen et al., 2021; Goddard et al., 2018).

## Data availability

All data is contained within the article.

## Supporting information

This article contains Supporting Information.

## Acknowledgements

We are grateful for Dr. David Vocadlo for helpful discussions and the kind gift of pharmacological inhibitors of OGT used in initial experiments for this study.

## Author contributions

Conceptualization: Sadia Rahmani, Warren W. Wakarchuk, Costin N. Antonescu Funding Acquisition: Warren W. Wakarchuk, Costin N. Antonescu Investigation: Sadia Rahmani, Hafsa Ahmed, Osemudiamen Ibazebo, Eden Fussner-Dupas. Formal Analysis: Sadia Rahmani, Hafsa Ahmed, Osemudiamen Ibazebo, Eden Fussner-Dupas, Costin N. Antonescu Supervision: Eden Fussner-Dupas, Warren W. Wakarchuk, Costin N. Antonescu Writing – original draft: Sadia Rahmani, Costin N. Antonescu Writing – Review & editing: Sadia Rahmani, Hafsa Ahmed, Osemudiamen Ibazebo, Eden Fussner-Dupas, Warren W. Wakarchuk, Costin N. Antonescu

## Funding and additional information

This work was supported by a Discovery Grant from the Natural Sciences and Engineering Research Council (RGPIN-2016-04371) and an Early Researcher Award from the Ontario Ministry of Research, Innovation and Science to C.N.A. S.R. was supported by an Ontario Graduate Scholarship.

