## Supporting Information for "O-GlcNAc transferase modulates formation of clathrin-coated pits"

**Supplementary Information Contents:**

**Table S1**: Antibodies and Chemicals

**Figure S1**: Overlay of AlphaFold generated AAK1 structures with experimentally resolved structures of AAK1
**Figure S2**: AAK1 residues predicted to be modified by O-GlcNAc and mutagenesis to generate O-GlcNAc deficient AAK1 mutant
**Figure S3**: Knockdown and rescue of AAK1
**Figure S4**: AAK1 WT and mutant display similar structure disorder prediction.
**Figure S5**: Slight overexpression of wild-type and GlcNAc deficient mutant of AAK1 similarly increase eGFP-clathrin intensity within clathrin labelled structures
**Figure S6**: Validation of CALM antibodies
**Figure S7**: LP-935509 treatment inhibits AP2 phosphorylation

**Table S1: Antibodies and Chemicals detailed information.**

| Name | Company | Product code |
|---|---|---|
| **Antibodies** | | |
| OGT/O-Linked-N-Acetylglucosamine Transferase | Abcam | ab177941 |
| O-GlcNAc RL2 | Abcam | |
| Anti-Phosphatidylinositol-binding clathrin assembly protein (Anti-PICALM) | Sigma Aldrich | HPA019061 |
| AAK1 | Cell Signaling Technology | 79832 |
| Phospho-AP2M1 (Thr156) | Cell Signaling Technology | #3843 |
| Epsin | Abcam | ab75879 |
| AP50 (AP2M1) | ThermoFisher Scientific | 27355-1-AP |
| clathrin heavy chain antibody [X22] | Abcam | Ab2731 |
| GAPDH | Cell Signaling Technology | 14C10 #2118 |
| Pan-Actin | Cell Signaling Technology | D18C11 #8456 |
| **Chemicals** | | |
| Thiamet G | Sigma Aldrich | SML0244 (Batch 22957) |
| LP-935509 (LP) | Axon MedChem | CAS [1454555-29-3] MF C20H24N6O3 |
| D-(+)-Glucose solution 45% in H2O, sterile-filtered, BioXtra, suitable for cell culture | Sigma Aldrich | G8769 |
| L-Glutamine BioXtra, suitable for cell culture | Sigma Alrich | G7513 |

| | | |
|---|---|---|
| DMEM, no glucose, no glutamine, no phenol red | ThermoFisher Scientific | A1443001 |

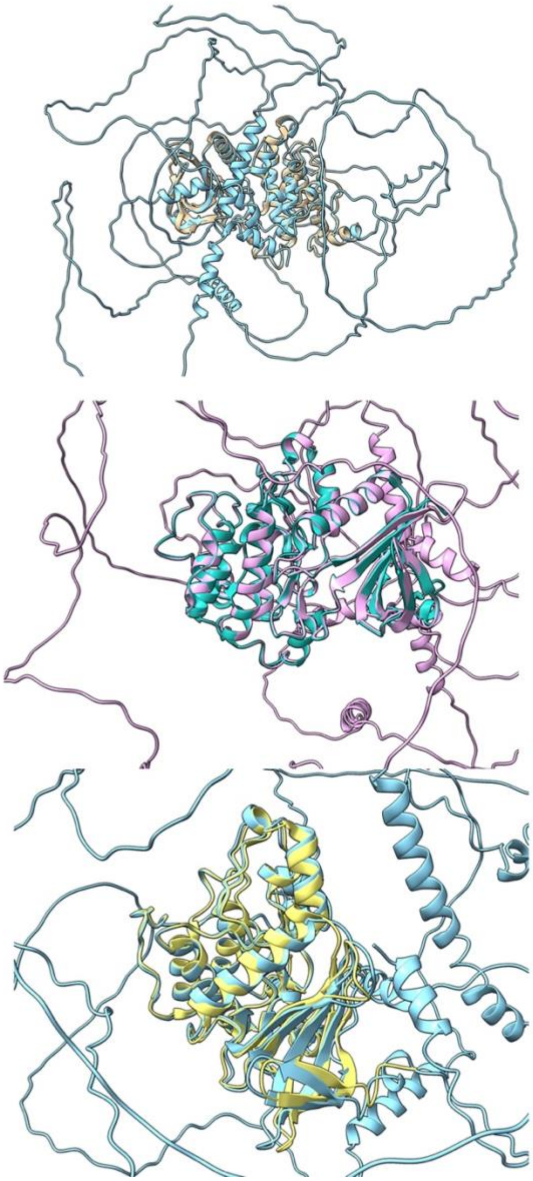

**Figure S1. Overlay of AlphaFold generated AAK1 structures with experimentally resolved structures of AAK1**. AAK1 wild type structure resolved by AlphaFold is compared to the experimentally resolved AAK1 structures: 4WSQ (PDB DOI: 10.2210/pdb4WSQ/pdb) (top panel), 5TE0 (PDB DOI: 10.2210/pdb5TE0/pdb) (middle panel), and 5L4Q (PDB DOI: 10.2210/pdb5L4Q/pdb) (bottom panel). The root mean squared deviation (RMSD) values for each 4WSQ, 5TE0, and 5L4Q as follows 0.557 Angstroms, 0.439 Angstroms, and 0.794 Angstroms respectively.

A

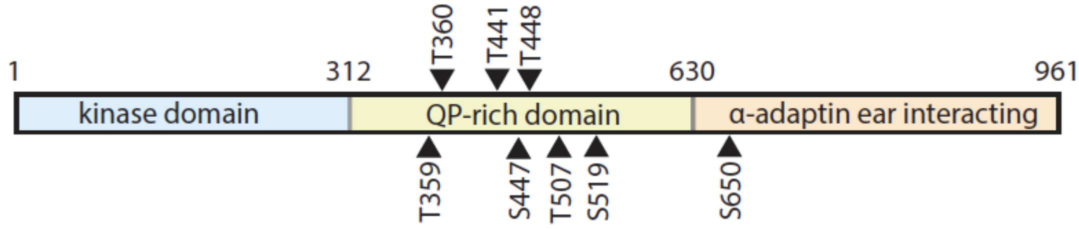


B

```
WT     RPDIYQVSYFSFKLLKKECPIPNVQNSPIPAKLPEPVKASEAAAKKTQPKARLTDPIPTT   360
MUT    RPDIYQVSYFSFKLLKKECPIPNVQNSPIPAKLPEPVKASEAAAKKTQPKARLTDPIPAA   360

WT     ETSIAPRQRPKAGQTQPNPGILPIQPALTPRKRATVQPPPQAAGSSNQPGLLASVPQPKP   420
MUT    ETSIAPRQRPKAGQTQPNPGILPIQPALTPRKRATVQPPPQAAGSSNQPGLLASVPQPKP   420

WT     QAPPSQPLPQTQAKQPQAPPTPQQTPSTQAQGLPAQAQATPQHQQQLFLKQQQQQQQPPP   480
MUT    QAPPSQPLPQTQAKQPQAPPAPQQTPAAQAQGLPAQAQATPQHQQQLFLKQQQQQQQPPP   480

WT     AQQQPAGTFYQQQQAQTQQFQAVHPATQQPAIAQFPVVSQGGSQQQLMQNFYQQQQQQQQ   540
MUT    AQQQPAGTFYQQQQAQTQQFQAVHPAAQQPAIAQFPVVAQGGSQQQLMQNFYQQQQQQQQ   540

WT     QQQQQQLATALHQQQLMTQQAALQQKPTMAAGQQPQPQPAAAPQPAPAQEPAIQAPVRQQ   600
MUT    QQQQQQLATALHQQQLMTQQAALQQKPTMAAGQQPQPQPAAAPQPAPAQEPAIQAPVRQQ   600

WT     PKVQTTPPPAVQGQKVGSLTPPSSPKTQRAGHRRILSDVTHSAVFGVPASKSTQLLQAAA   660
MUT    PKVQTTPPPAVQGQKVGSLTPPSSPKTQRAGHRRILSDVTHSAVFGVPAAKSTQLLQAAA   660
```

**Figure S2. AAK1 residues predicted to be modified by O-GlcNAc and mutagenesis to generate O-GlcNAc deficient AAK1 mutant.** (A) The domain structure of AAK1, a 961 amino acid protein that has three main domains: Kinase domain, QP rich domain and AP2 interaction domain. The predicted O-GlcNAc sites fall mainly in the QP rich domain. (B) We identified O-GlcNAc sites on AAK1, and mutated them to Ala: T359A, T360A, T441A, S447A, T448A, T507A, S519A, S650A.

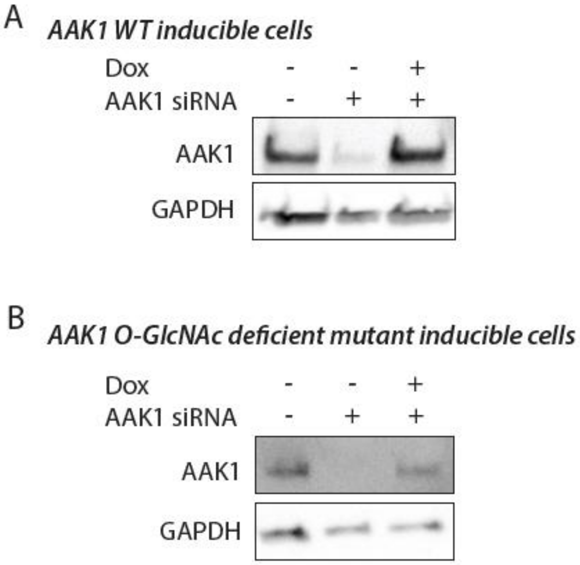


**Figure S3. Knockdown and rescue of AAK1.** ARPE-19 cells were engineered to stably harbor a transgene for inducible expression of eGFP-tagged AAK1 wild-type (WT) or harboring mutations in the 8 predicted O-GlcNAc modification sites (mut). Cells were treated with siRNA to silence endogenous AAK1 as shown. Following siRNA transfection and treatment with dox for 24h, whole cell lysates were resolved by western blotting. Shown are representative western blots using antibodies to detect AAK1 expression or GAPDH (loading control). This strategy achieves a slight overexpression of wild-type AAK1 and a near endogenous level of expression of mutant AAK1. The migration of the eGFP-tagged AAK1 (~165kDa) is similar to that of endogenous AAK1 (~145kDa) under these western blotting conditions.

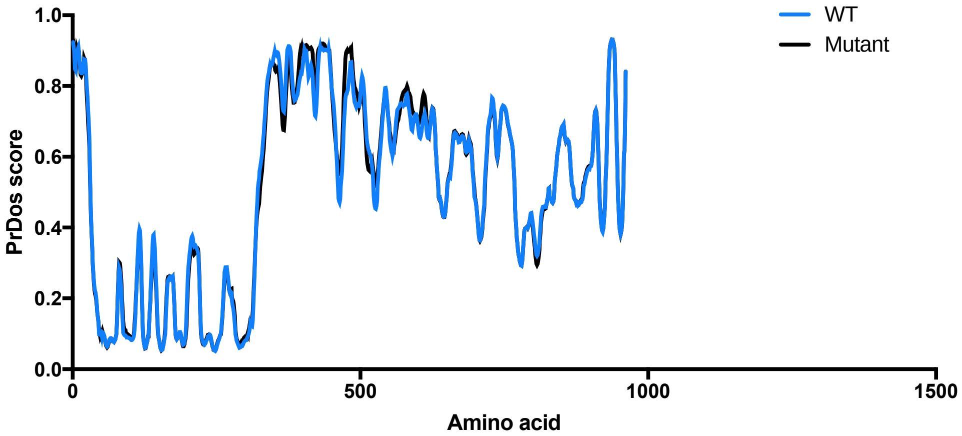


**Figure S4: AAK1 WT and mutant display similar structure disorder prediction.** Protein Disorder was determined using Protein Disorder Prediction System serve (PrDOS) using the default settings. A scatter plot was generated using the disorder predictions of both the WT and the mutant forms of AAK1.

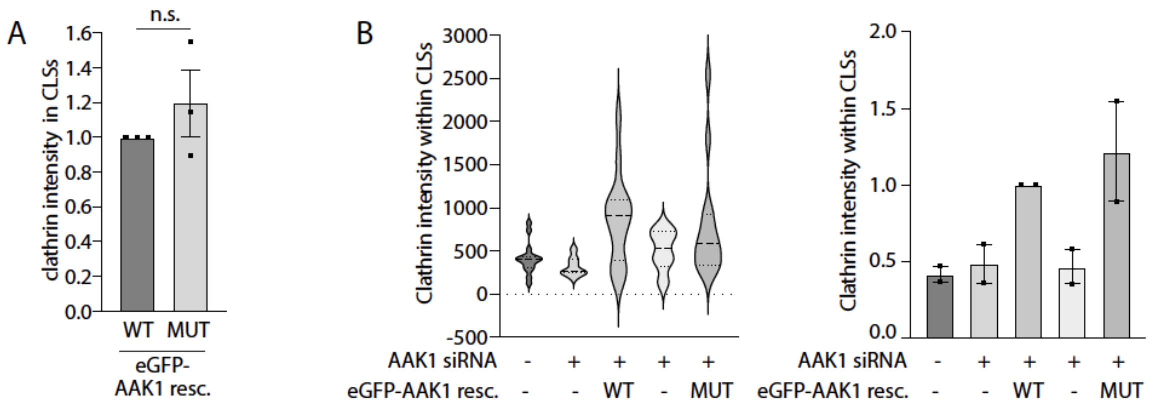


**Figure S5. Slight overexpression of wild-type and GlcNAc deficient mutant of AAK1 similarly increase eGFP-clathrin intensity within clathrin labelled structures**. ARPE-19 cells were engineered to stably harbor a transgene for inducible expression of eGFP-tagged AAK1 wild-type (WT) or harboring mutations in the 8 predicted O-GlcNAc modification sites (mut). Cells were treated with siRNA to silence endogenous AAK1, or non-targeting siRNA (control). Following siRNA transfection and treatment with dox, cells were fixed and stained to label clathrin heavy chain, followed by imaging by TIRF and epifluorescence microscopy. Experiment is as described in Figure 4D-E, and representative images are shown in Figure 4D. Images were subjected automated detection and analysis of CLSs as described in *Materials and Methods*. (A) Shown are the measurements of clathrin intensity within CLSs comparing only the rescue conditions (with either eGFP-AAK1 wild type and mutant) from *n*=3 independent experiments. (B) Shown are the measurements of clathrin intensity as the distribution of these values in a representative experiment as a violin plot (left panel), or the mean (bar) ± SE from 2 independent experiments, as well as individual values from each experiment (dots).

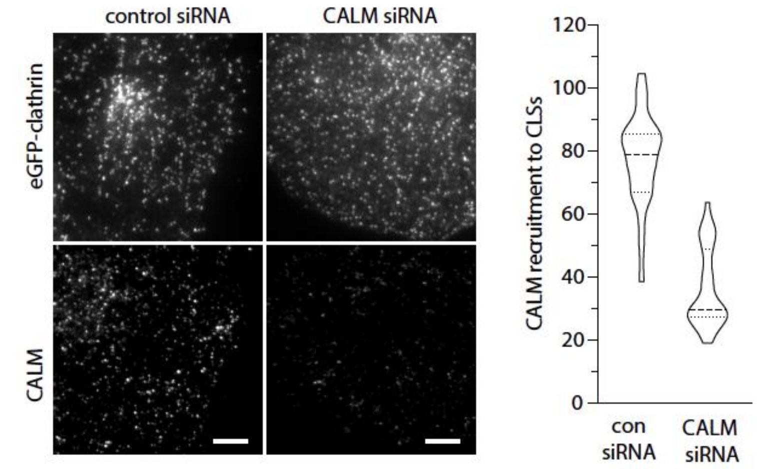


**Figure S6. Validation of CALM antibodies for immunofluorescence detection using TIRF microscopy.** ARPE-19 cells were treated with siRNA targeting CALM or with non-targeting siRNA (control, con). ARPE-19 cells stably expressing eGFP-clathrin light chain were treated with siRNA to silence CALM or non-targeting (control) siRNA. Following transfection, cells were fixed and stained with anti-CALM antibodies, followed by imaging by TIRF microscopy, with representative images shown in the left panels. Scale 5 µm. CLSs were subjected to automated detection and analysis as described in *Materials and Methods*. Shown are the distributions as a violin plot of measurements of eGFP-clathrin in individual CLSs within a representative experiment. This indicates the specificity of the fluorescence signal in TIRF microscopy derived from the antibodies used to detect endogenous CALM by immunofluorescence staining.

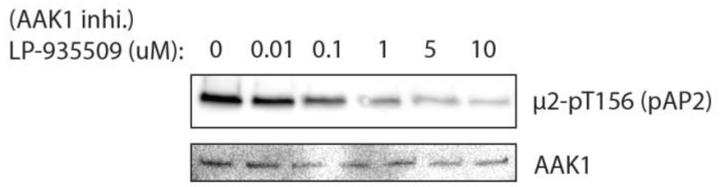


**Figure S7: LP-935509 treatment inhibits AP2 phosphorylation.** ARPE-19 cells were incubated with different concentrations of LP-935509 as shown for 3 h before lysate preparation. Shown is a representative western blot using whole-cell lysates, showing detection of pAP2 (phospho-Thr156 on the µ2 subunit of AP2) or AAK1.
